# Electrocommunication signals indicate motivation to compete during dyadic interactions of an electric fish

**DOI:** 10.1101/2021.02.04.429572

**Authors:** Till Raab, Sercan Bayezit, Saskia Erdle, Jan Benda

## Abstract

Animals across species compete for limited resources. While in some species competition behavior is solely based on own abilities, others assess their opponents to facilitate these interactions. Using cues and communication signals, contestants gather information about their opponent, adjust their behavior accordingly, and can thereby avoid high costs of escalating fights. We tracked electrocommunication signals, in particular “rises”, and agonistic behaviors of the gymnotiform electric fish *Apteronotus leptorhynchus* in staged competition experiments. A larger body size relative to the opponent was the sole significant predictor for winners. Sex and the frequency of the continuously emitted electric field were only mildly influencing competition outcome. In males, correlations of body-size and winning were stronger than in females and, especially when losing against females, communication and agonistic interactions were enhanced, hinting towards males being more motivated to compete. Fish that lost competitions emitted the majority of rises in dependence on the competitors’ relative size and sex. The emission of rises was costly since it provoked ritualized biting or chasing behaviors by the other fish. Despite winners being accurately predictable based on rise numbers already after the initial 25 minutes, losers continued to emit rises. The number of rises emitted by losers and the duration of chasing behaviors depended in similar ways on physical attributes of contestants. The detailed evaluation of these correlations hint towards *A. leptorhynchus* adjusting their competition behavior according to mutual assessment, where rises could signal a loser’s motivation to continue assessment through ritualized fighting.

## Introduction

Across animal species fighting is a key behavior to secure access to limited resources (Clutton-Brock et al., 1979; Chapman et al., 1995; Markham et al., 2015). However, competition is costly because of the energy and time allocated to it and the increased risk of injury or death (e.g. Briffa and Elwood, 2004). Therefore, individual behavioral decisions during contests are strongly dependent on the associated potential benefits and costs (Arnott and Elwood, 2008, 2009). Often, the best predictor for the outcome of competitions is the contestant’s fighting ability, also called resource holding potential (RHP, Parker, 1974). Usually, larger and stronger individuals win contests, because their physical advantages, i.e. higher RHP, directly reflect their increased endurance and potential to inflict damage during contests (Archer, 1988). Additional factors like weaponry, experience, sex, or positional advantages also influence RHP (reviewed in Arnott and Elwood, 2008).

Behaviors and the course of a competition have been shown to be either based on the assessment of solely the own RHP or by integrating both, the own and opponent’s RHP (Taylor et al., 2001; Davies and Halliday, 1978; Enquist et al., 1990; Huyghe et al., 2005). In the first case (self-assessment), costs resulting from competition are accumulated until an endurance threshold, set by an individual’s RHP, is reached and the respective individual retreats (Arnott and Elwood, 2009). Competition costs either arise exclusively from own behaviors (pure self-assessment, Taylor and Elwood, 2003) or are supplemented by costs inflicted by opponents (cumulative assessment, Payne, 1998). In both cases no direct information about an opponent and its RHP is gathered. Alternatively, in mutual assessment the contestants assess each other’s RHP, compare it to the own, and adjust their behavior according to the difference (Enquist and Leimar, 1987). The huge benefit of this strategy is its economic efficiency. Individuals can recognize their inferiority and retreat long before their endurance threshold is reached, thereby saving metabolic costs for both competitors.

Besides passive cues signaling RHP, actively produced communication signals may facilitate interactions during animal conflict (Arnott and Elwood, 2009; Seyfarth et al., 2010). They can directly indicate and reflect an individual’s RHP (Davies and Halliday, 1978; Clutton-Brock et al., 1979), but also convey information about additional factors influencing a contest and its outcome, like motivation and behavioral intent (e.g. aggression: Kareklas et al., 2019; Westby and Box, 1970, or submission: Hupé and Lewis, 2008; Batista et al., 2012) or social status (Fernald, 2014). Such low cost signals have been shown to reduce the intensity and duration of contests or even convey sufficient information to resolve conflicts without the necessity of physical competitions (Parker, 1974; Clutton-Brock et al., 1979; Janson, 1990).

To prevent the high costs of repetitive fights with the same oppponents, dominance hierarchies are established in various species (Creel et al., 1996; Janson, 1985; Clutton-Brock et al., 1979). In domoninace hierarchies the necessity of fighting is reduced since access to resources is regulated through social status, favoring those individuals of higher rank (Janson, 1985; Wauters and Dhondt, 1992; Sapolsky, 2005; Taves et al., 2009). The organization and characteristics of dominance hierarchies vary across species (Janson, 1985; Cigliano, 1993; Sapolsky, 2005). While in group-living species complex social structures, like for example a leader-follower dynamic, can emerge (Strandburg-Peshkin et al., 2018), in solitary species dominance is rather associated with resource-based benefits, like the occupation of higher quality territories and increased reproductive success (e.g. Cigliano, 1993). Differences in the abundance and dispersion of food can further lead to variations regarding the skewness in access to resources across social ranks. In bottom-up egalitarian hierarchies resources are more equally distributed (Sapolsky, 2005), whereas in top-down despotic hierarchies access to resources is strongly skewed in favor for dominant individuals (Kappeler and Schäffler, 2008).

Dominance hierarchies have also been suggested for the nocturnal gymnotiform electric fish *Apteronotus leptorhynchus* (Dunlap and Oliveri, 2002; Stamper et al., 2010; Raab et al., 2019). *A. leptorhynchus* competes for mates during mating season only (Hagedorn and Heiligenberg, 1985; Henninger et al., 2018), otherwise they rival for optimal shelters (Dunlap and Oliveri, 2002). The corresponding competitions are characterized by ritualized fighting behaviors accompanied by electrocommunication signals (Triefenbach and Zakon, 2008; Smith, 2013). While body-size has been shown to be the main determinant for the outcome of competitions in gymnotiformes (Batista et al., 2012; Triefenbach and Zakon, 2008; Dunlap and Oliveri, 2002), the influence of other factors like sex and communication signals require further investigation.

Electric signaling has been shown to be an integral aspect of agonistic behaviors in gymnotiform fish (Westby and Box, 1970; Batista et al., 2012; Triefenbach and Zakon, 2008; Hupé and Lewis, 2008; Henninger et al., 2018). Already the frequency of their continuous electric organ discharge (EOD) has been suggested to signal an individual’s physical condition, dominance status, or aggressiveness (Westby and Box, 1970; Hagedorn and Heiligenberg, 1985; Cuddy et al., 2012). The sexually dimorphic EOD frequency (EODf) of *A. leptorhynchus* indicates identity and sex (Henninger et al., 2020), with males having higher EODfs than females (Meyer et al., 1987). While some studies also suggest EODf to indicate dominance (Hagedorn and Heiligenberg, 1985; Dunlap and Oliveri, 2002; Cuddy et al., 2012; Henninger et al., 2018; Raab et al., 2019), others were not able to replicate this correlation (Triefenbach and Zakon, 2008). For generating distinct electrocommunication signals electric fish modulate their EODf on various time scales (Benda, 2020). One category of electrocommunication signals, so called “rises”, are characterized by a moderate increase in EODf by no more than a few tens of Hertz followed by an approximately exponential decay back to baseline EODf over a second to almost a minute (Hopkins, 1974; Engler et al., 2000; Tallarovic and Zakon, 2002; Zakon et al., 2002). The function of rises is still discussed controversially. They have been suggested to signal aggression or motivation to attack (Tallarovic and Zakon, 2005; Triefenbach and Zakon, 2008), submission (Hopkins, 1974; Serrano-Fernández, 2003), “victory cries” (Dye, 1987), to evoke or precede attacks (Hopkins, 1974; Triefenbach and Zakon, 2008), or to just by a general expression of stress (Smith, 2013). An enhancement of sensory acquisition by rises is highly unlikely, because their mild increase in EODf only marginally influences encoding in electrorecptor afferents (Walz et al., 2014).

Using recently developed techniques for tracking electrocommunication signals in freely behaving electric fish (Henninger et al., 2018; Madhav et al., 2018; Henninger et al., 2020) in addition with infrared video recordings, we recorded electric and physical interactions of pairs of *A. leptorhynchus* in staged competitions over a superior shelter. Compared to previous studies we significantly expanded the observation times (from ten minutes to six hours) and the number of interacting pairs of fish. We evaluated the influence of body size, weight, sex, and EODf on the outcome of competitions. Analyzing relations between rises, agonistic interactions, and physical difference between competitors we were able to uncover the fish’s assessment strategy, quantify behavioral difference between the sexes, and identify potential uses of rises of *A. leptorhynchus* during competitions.

## Methods

### Aminals

A total of 21 mature *A. leptorhynchus* (9 males, 12 females) not in breeding condition, obtained from a tropical fish supplier, were used. Fish were selected randomly from multiple populations to reduce familiarity effects and sorted into four mixed-sex groups of five or six fish (males/females in group 1: 2/4, 2: 1/4, 3: 3/2, 4: 3/2). Males were identified by their higher EODf (see below) and elongated snout. The sex of half the fish was verified after the competition experiments via post mortem gonadal inspection in the context of electrophysiological experiments (approved by Regierungspräsidium Tübingen, permit no. ZP 1/16). Fish were housed individually in 54 l tanks at a light-dark cycle of 12 h/12 h prior to the experiments and in between competition trials. Each tank was equipped with a plastic tube for shelter, a water filter, and an electrical heater. Water temperature was constant at 25 ± 0.5 °C and water conductivity 200 µS*/*cm. Fish were fed frozen *Chironomus plumosus* on a daily basis. The competition experiments complied with national and European law and were approved by the Regierungspräsidium Tübingen (permit no. ZP 04/20 G).

### Setup

The competition experiments were run in a 100 l tank equipped with a 10 cm long and 4 cm wide PVC half-tube as a superior shelter in the center, surrounded by four additional, less optimal shelters (two 5 cm long, 4 cm diameter PVC half-tubes and two 3 × 5 cm^2^ tables, fig. S-1). Water temperature and conductivity as well as light-dark cycle were identical to those in the housing tanks. A heating mat was placed below the tank and powered with DC current. Two air-powered water-filters were placed behind PVC boards with netted windows in the corners of the tank to not offer additional shelter. 15 monopolar electrodes at low-noise buffer headstages were mounted on the bottom of the tank. The reference electrode was placed behind a PVC board in one corner of the tank. Electric signals were amplified (100 × gain, 100 Hz low-pass filter, EXT-16B, npi electronic, Tamm, Germany) digitized at 20 kHz per channel with 16 bit resolution (USB-1608GX-2AO, Measurement Computing) and stored on 64 GB USB sticks using custom written software running on a Raspberry PI 3B. Water temperature was measured every 5 min (Dallas DS18B20 1-wire temperature sensor). Infrared-videos were recorded at 25 frames per second with a camera (Allied Vision Guppy PRO) mounted on top of the tank for all trials of groups 3 and 4. The tank was continuously illuminated by 2 × 4 infrared lights (ABUS 840 nm) located on the long sides outside the tank. For the synchronization of video and electric recordings we used LED-light pulses of 100 ms duration triggered by the computer-amplifier system in intervals of 5 seconds. The LED was mounted on the edge of the tank not perceivable by the competing fish, but detectable in the video-recordings. The tank, camera and illumination was placed inside a faraday cage.

### Experimental procedure

In each competition trial two fish were freely swimming and interacting in the experimental tank for 6 h. Participating fish were taken from their housing tanks and simultaneous released into the experimental tank. The first 3 h of each trial took place during the dark-phase and the second 3 h during the light-phase of the circadian rhythm the fish were accustomed to. This limited the experiment to one trial per day. The winner of each trial was identified by its presence within the superior shelter during the light-phase of the trial. Fish were transferred back into their housing tanks after the trial. Pairings for each trial were selected systematically to (i) ensure all possible combinations within each group to be tested (10 combinations for groups of five fish, 15 for the six-fish group), (ii) keep the experience level for all fish equal, and (iii) prevent a single fish to be tested on two consecutive days. Weight and length (body size) of each fish was assessed once a week starting in the week before the competition trials.

With 21 fish in four groups we ended up with a total of 45 pairings and trials. Technical failure lead to loss of the electric recordings for the initial four trials of group four. In another three trials we were unable to extract EODf traces and electrocommunication signals from the electric recordings because the EODf difference between fish were to low (< 0.5 Hz). In a single trial, that we discuss separately, we were unable to determine the winner, because both fish shared the superior shelter at the end of the trail. The remaining 37 trials were analyzed in detail.

### Preprocessing of electric and video recordings

After computing spectrograms for each channel with fast Fourier transformation (FFT, *n*_fft_ = 2^15^, corresponds to 1.63 s, 80 % overlap) we first detected peaks in a power spectrum summed over the channels and assigned them to the fundamental EODfs and their harmonics of the two fish. Based on EODfs and the distribution of power in the channels we tracked electric signals over time and obtained EODf traces for each of the two fish (Henninger et al., 2018, 2020; Madhav et al., 2018).

To asses baseline EODf we computed the 5th percentile of non-overlapping, 5 minute long EODf trace snippets. EODf is sensitive to temperature changes, which were inevitable throughout the single trials and averaged at 1 °C. We computed the *Q*_10_-values resulting from temperature and EODf differences of each 5 minute snippet and used the median of 1.37 over all fish to adjust each fish’s EODf to 25 °C (EODf_25_). The EODf_25_ was used to assess an individuals sex, with EODf_25_ > 740 Hz assumed to originate from males. As noted above, the sex of half of the fish was verified via post mortem gonadal inspection.

EODf difference for each pair of fish was estimated from the difference of the median baseline EODfs of the competitors during the light-phase of each trial, where EODfs stayed comparably stable. Rises were identified by detecting their characteristic onset peak in each EODf trace based on a minimum difference of 5 Hz between a peak and the preceding trough (Todd and Andrews, 1999). Then the size of the rise, its maximum increase in EODf, was calculated by subtracting the baseline EODf from this peak frequency.

We manually extracted two categories of agonistic interactions from the infrared video recordings using the event logging software BORIS (Friard and Gamba, 2016). For agonistic interactions without physical contact that were characterized as high velocity, directed movements towards a competitor (e.g. chasing behavior) we recorded onset and end times. Agonstic physical contacts between competitors like ritualized biting or head bumping were detected as point events.

### Data analysis

Data were analyzed in Python version 3.6.8 using numpy (van der Walt, 2011), scipy (Oliphant, 2007) and matplotlib (Hunter, 2007) packages . All averages are given with their standard deviation. Mann-Whitney *U* tests were used to asses significance of differences between two populations and Pearson’s test and correlation coefficient *r* for assessing correlations. The influence of various factors on competition outcome was quantified by paired *t*-tests and by the area under the curve (AUC) of a receiver-operating characteristics (ROC). Generalized linear mixed models (GLM)

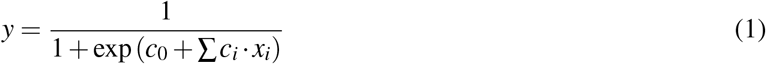

with a logistic link function were used to estimate the combined effects of several factors *x*_*i*_ (continuous: EODf, size, ΔEODf and Δsize, categorical: sex), linearly combined with coefficients *c*_*i*_ and an offset *c*_0_, on the outcome of the competitions *y* (winner or looser). The performance of the GLMs was again assessed by the AUC of a ROC-analysis. Standard deviations of AUC values were obtained by 1000 times bootstrapping the data.

To evaluate the influence of the contestant’s physical attributed on the quantity of emitted rises throughout a trial simple correlations were supplemented by multiple linear regression models. For each model we performed backwards elimination model selection with an elimination criterion of α > 0.05.

Temporal coupling between rises and agonistic interactions was quantified by a cross-correlation analysis, i.e. by estimating the averaged rate of rises centered on agonistic events (onset of chasing behaviors or physical contact). For this the temporal differences, Δtime, between agonistic onsets and rises up to ±60 sec were convolved with a Gaussian kernel with a standard deviation of 1 s. Statistical significance was assessed by a permutation test. The null hypothesis of rises not being correlated with agonistic interaction events was obtained by computing the cross-correlation as described above from 1000 random variants of shuffled rise intervals. From this distribution we determined the 1st and 99th percentiles. In addition, we computed the 98 % confidence interval for the estimated cross-correlation by 1000 times jackknife resampling where we randomly excluded 10 % of the rises.

We used a time window 5 s prior to agonistic onsets to quantify the average number of rises per agonistic event and to compare them to the corresponding time fractions, the number of agonistic events multiplied with the 5 s window relative to the total dark-period time of 3 h . The time window of 5 s was chosen to approximately cover the time of significantly elevated rise rates that we observed before the agonistic events in the cross-correlation analysis.

## Results

In 37 trials we observed and analyzed pairs of *A. leptorhynchus* (6 male pairs, 10 female pairs, and 21 mixed-sex pairs) competing for a superior shelter. The 9 males differed from the 12 females by their higher EOD frequency (EODf) as expected from the sexual dimorphism in EODf in *A. leptorhynchus*. The fish’s size ranged from 9 to 19 cm and was independent of sex (*U* = 51.5, *p* = 0.443, fig. S-1 A). Fish size strongly correlated with body weight (3.3–20.3 g, *r* = 0.94, *p* < 0.001, fig. S-1 B) and we therefore excluded weight from the following analysis.

We were able to track electric behaviors of the competitors based on the individual-specific EODf traces, including the detection of rises (electrocommunication signals, fig. 1 A,B). Complementary infrared-video recordings obtained during the 20 trials of group 3 and 4 were used to detect agonistic behaviors, i.e. chasings (fig. 1 C) and physical contacts like ritualized biting or head bumping (fig. 1 D). In a typical competition trial (fig. 1 E) the overall activity of the two fish was much higher during the initial, three hour dark phase as demonstrated by the high rates of chasing and contact events as well as of rises. During the subsequent three hour light phase, the activity ceased almost entirely and one fish spent substantially more time within or closer to the superior shelter (99.27 ± 0.006 %). This fish was identified as the winner of the competition trial. EODfs of the two fish usually differed clearly and decreased over the course of the experiment because of slightly decreasing water temperature.

**Figure 1:**
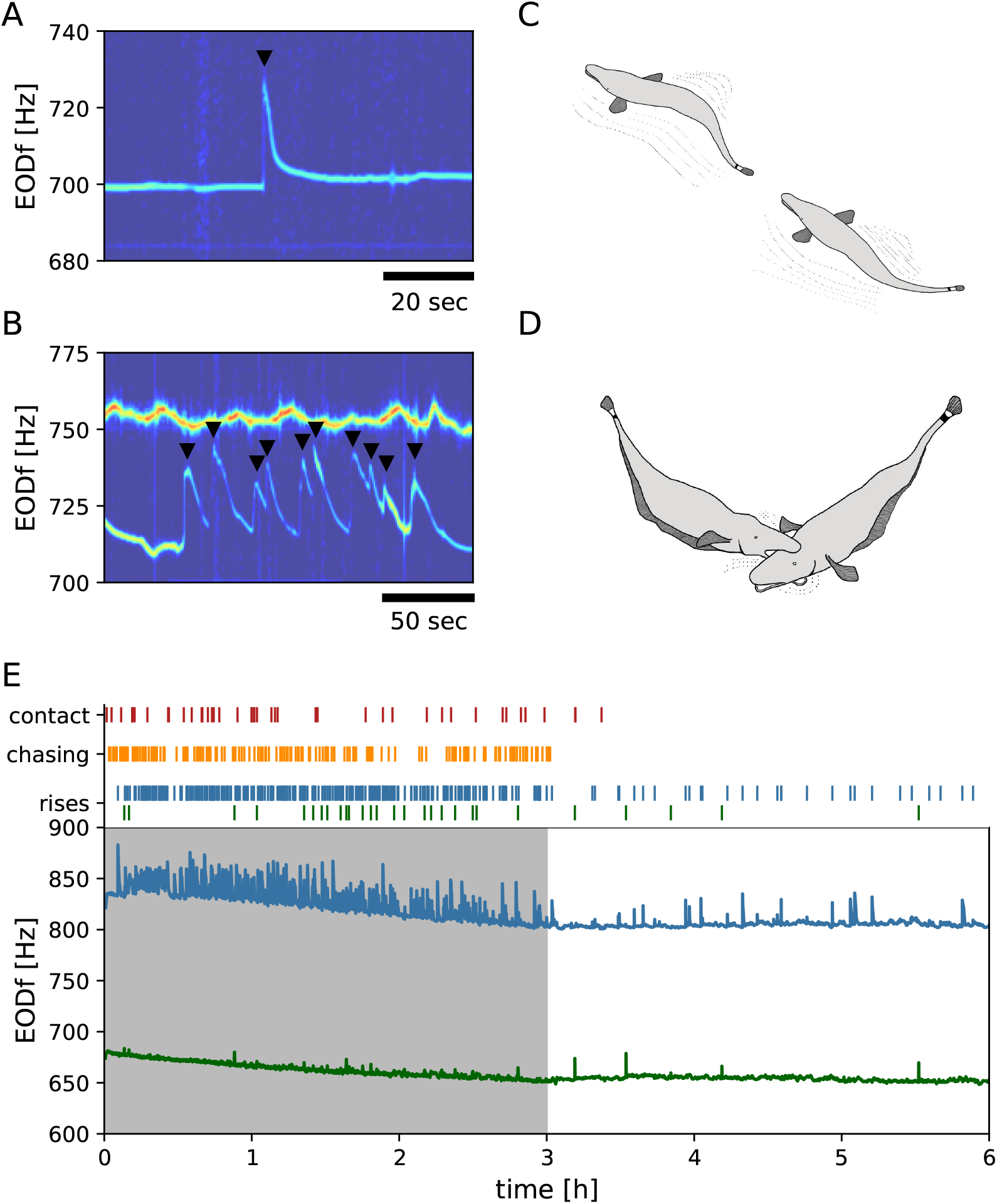
Behaviors and interactions of *A. leptorhynchus* during a typical competition trial. **A** Spectrogram of an electric recording including the EODf trace of a single fish while emitting a rise as communication signal. Rises are abrupt increases in EODf followed by an exponential decay back to baseline EODf. Rises were detected using their characteristic onset peak (black triangle). **B** Spectrogram of a 200 second snipped of a electric recording comprising EODf traces of two fish. In the lower EODf trace a series of 10 rises can be seen. **C, D** Ritualized agonistic interactions in *A. leptorhynchus* comprise non-physical, high velocity chasing events (panel C) and short physical agonistic behaviors like biting or dead bumping (panel D). Both were initiated by fish later winning a trial. **E** *A. leptorhynchus* continuously emits EODs with an individual specific frequency (EODf). EODf traces of both competing fish (blue male and green female, bottom panel), time points of physical contacts, onsets of chasing behavior, and detected rises (top panels) recorded during the full 6 h trial with the first 3 h in darkness (gray background) and the last 3 h during light.

### Larger fish win competition

Larger fish were more likely to win the superior shelter (*t* = 5.3, *p* < 0.001, fig. 2 A). Over all pairings, winners are correctly assigned with a probability of 90 % based on size difference (in the sense of the AUC of a ROC analysis, fig. S-3 C). In particular in trials won by males, most of the winners were larger than the losers (male-male: 5 out of 6, *t* = 2.9, *p* = 0.04; male-female: 11 out of 14, *t* = 3.9, *p* = 0.002; fig. 2 B, D). In trials won by females, this influence of size difference was similarly pronounced but not significant (female-female: 8 out of 10 winners were larger, *t* = 2.1, *p* = 0.07; female-male: 6 out of 7 winners were larger, *t* = 2.0, *p* = 0.09; fig. 2 C, E). In 12 of the 21 mixed-sex pairings were the males larger than the competing females, of which only a single male lost. Of the nine larger females three lost. Absolute size, on the other hand, did not predict competition outcome (AUC = 67 %, fig. 2 F). Note that neither in males nor in females EODf correlated with size (males: *r* = 0.47, *p* = 0.20; females: *r* = 0.04, *p* = 0.90) and that size was independent of sex (fig. S-1 A).

**Figure 2:**
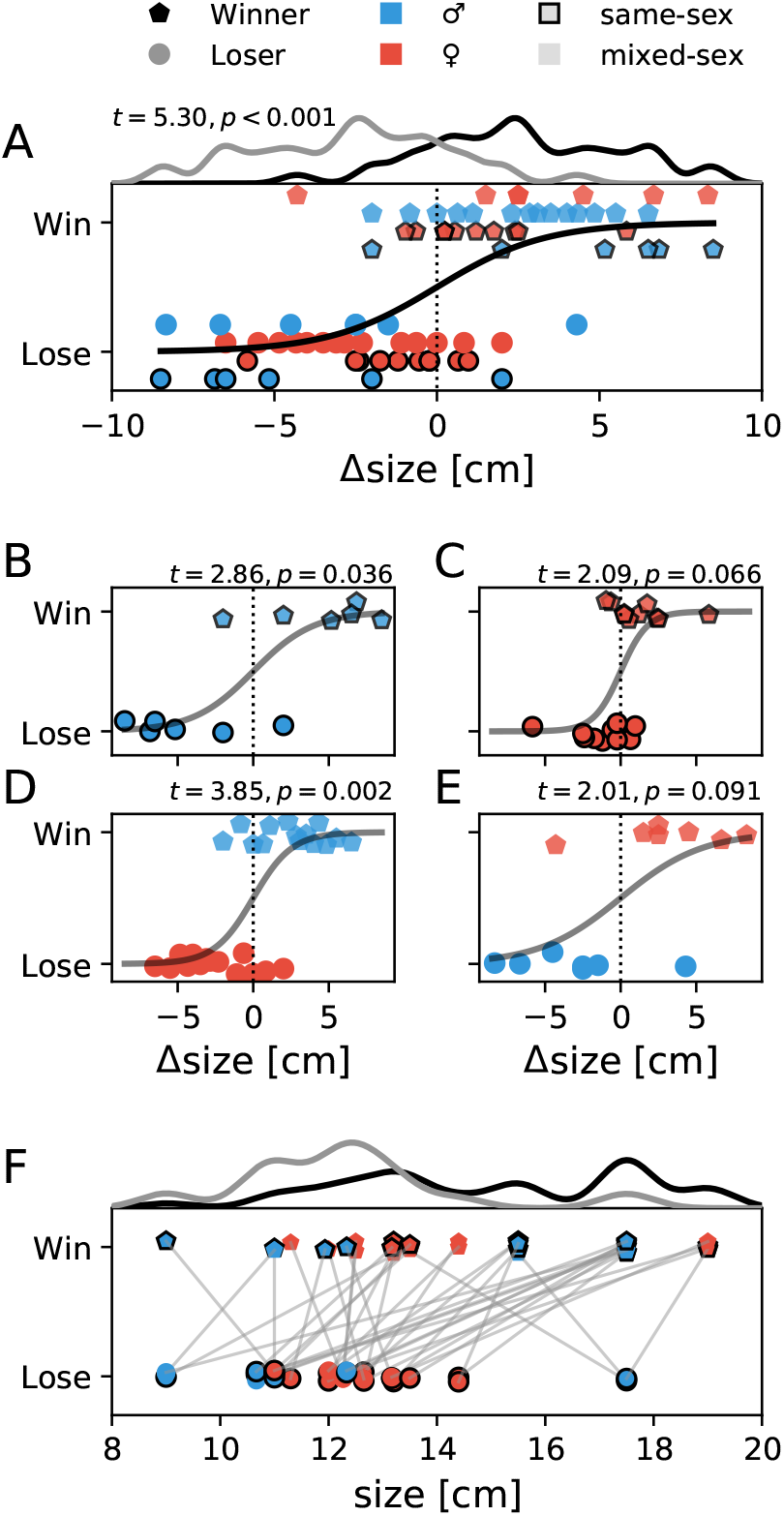
Body size of winners and losers. Colors and marker style indicate different pairings and the outcome of the competition. Each competition trial contributes two data points, one for the winner and one for the loser. Blue represents males, red females. Pentagons indicate winners, circles losers. Black marker edges indicate same-sex pairings. Winners and losers of each sex pairing were offset in panels A and B and jittered in panels C–F. **A** Winners were larger than their opponents as indicated by a logistic fit (black line, *t* = 5.3, *p* < 0.001) and kernel histograms of winners and losers (AUC = 90 %). Size differences are measured relative to the opponent. **B, C, D, E** Winners were larger in all sex pairings. In particular, winning males were larger than their male (B) or female (D) opponent. **F** Distributions of absolute body size of winners and losers largely overlap. The crucial factor determining winners and losers is the difference in size between opponents as shown in A. Grey lines connect pairs competing in a trial.

### Fish with higher EODf seem to win competition

Previous studies suggest EODf to be an indicator for dominance in *A. leptorhynchus* (Dunlap and Oliveri, 2002; Henninger et al., 2018; Raab et al., 2019). Indeed, winning fish had on average higher EODfs in comparison to their opponents (*t* = 2.1, *p* = 0.04, fig. S-2 A). However, in same sex pairings analyzed separately, EODf did not predict competition outcome neither in males (*t* = 0.79, *p* = 0.47, fig. S-2 B) nor in females (*t* = 1.4, *p* = 0.19, fig. S-2 C), but the positive coefficient of the logistic regressions suggests a mild influence of EODf on the outcome of competitions. Further, in mixed sex pairings, winning males always had higher EODfs and winning females always lower EODfs than their opponent, because of the sexual dimorphic EODfs in *A. leptorhynchus* (fig. S-2 D & E). These mixed-sex competitions were more often won by males than by females (14 out of 21, Binomial test 14 or more males winning assuming equal chances for both sexes: *p* = 0.10). This asymmetry results in an AUC = 82 % for discriminating winners from losers based solely on sex as a rough proxy for EODf (males have high, females low EODf). Although the difference in EODfs potentially contains more detailed information than sex alone, it does not discriminate winners from loser better than sex (AUC = 75 %). Absolute EODf is even less informative about competition outcome (AUC = 65 %, fig. S-2 F).

### Factors influencing competition outcome

We constructed a generalized linear mixed model (GLM, Eq. (1)), predicting the competition outcome based on all the measured physical factors size, size difference, EODf, EODf difference, and sex (fig. S-3 A–C) for estimating an upper bound of discrimination performance. As expected from the single-factor analysis, size difference is the only factor significantly contributing to the prediction of winners (*t* = 2.4, *p* = 0.02, tab. 1). The model correctly predicts the outcome of 34 of the 37 competition trials with an AUC of 94 % (fig. S-3 C). Two-factor GLMs based on size differences and either sex or EODf differences perform similarly well (AUC = 93 %) and slightly better than size difference alone (AUC = 90 %), further questioning the role of EODfs in predicting competition outcome (fig. S-3 C). The outcome of competitions were independent from previous encounters. Auto correlations of winlost-histories did not differ from those of random sequences, where winners and losers were assigned randomly (permutation test).

**Table 1:** Generalized linear mixed model assessing the significance of difference physical factors on winning competitions. For each factors *i* coefficients *c*_*i*_ (Eq. (1)), *t*-statistics, and significance is given. The size difference between competitors is the only significant factor of the model, indicating its predominant importance for winning competitions.

| factor | coefficient | t-value | p-value |
| --- | --- | --- | --- |
| sex<sub>f1</sub> | −1.437 | −0.498 | 0.618 |
| sex<sub>f2</sub> | 0.174 | 0.067 | 0.946 |
| size<sub>f1</sub> | 0.046 | 0.255 | 0.799 |
| size<sub>f2</sub> | −0.371 | −1.918 | 0.055 |
| EODf<sub>f1</sub> | 0.002 | 0.199 | 0.842 |
| EODf<sub>f2</sub> | −0.005 | −0.456 | 0.648 |
| Δsize | 0.417 | 2.392 | 0.017 |
| ΔEODf | 0.007 | 0.760 | 0.447 |

### Rises

We detected in total 8 530 rises using their characteristic onset peak in EODf (fig. 1 A & B). The size of the rises, the peak EODf relative to baseline EODf before, ranged from the detection threshold of 5 Hz up to 68 Hz with a mean of 17 Hz. We were not able to detect any dependency of our results on the size of rises. Rises categorized in distinct size categories did not behave differently. In the following we therefore focus on an analysis of rise numbers and timing.

### Losing fish emit more rises during the active phase

Rises were primarily emitted during the dark-phases, i.e. when fish were actively interacting with each other (*t* = 6.7, *p* < 0.001). The fish that later in the light phase did *not* occupy the superior shelter, produced ten-fold more rises in the dark phase (184 ± 105) than their winning competitors (18 ± 17, *t* = 9.5, *p* < 0.001, fig. 3 A). Loser rise counts were highly variable, they ranged from 0 to 419 rises per trial with a coefficient of variation of *CV* = 0.63.

**Figure 3:**
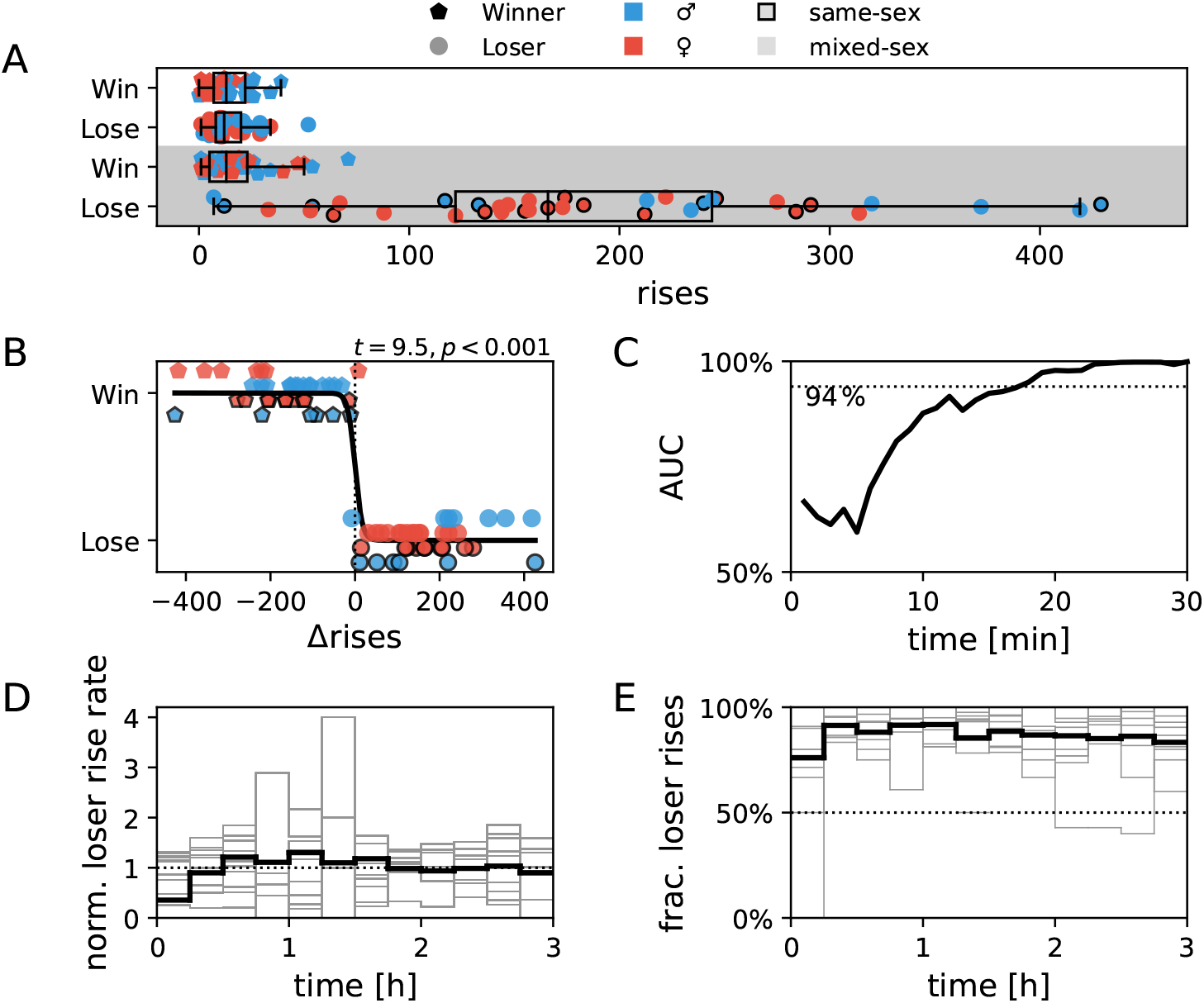
Rise rates of winners and losers. **A** Rises were predominantly produced by losers of competitions during the dark-phase (shaded). Winners during the dark-phase produced equally little rises as both winners and losers during the light-phase (white background). **B** Losers of competitions reliably produced more rises than winners in the dark, although absolute and relative numbers of rises varied considerably between trials. **C** Time resolved discrimination performance between winners and losers based on differences in cumulative rise counts for the first 30 min of the trials. The AUC of ROC analysis asymptotes to almost 100 % after about 25 min, indicating the outcome of the trials to be already determined the latest from this time on. For comparison, the dashed line at 94 % indicates the performance of the GLM including all physical characteristics of the fish from fig. S-3 C. **D** Time course of loser rise rates during the dark phase. Rise rates of each trial (gray) were normalized to their mean rate. On average (black) rise rates reached a constant level after about 30 min. **E** Fraction of rises of losing fish quantified in 15 min time windows is consistently larger than 50 % (dashed line) throughout the whole dark period. Individual trials in gray, average over trials in black.

The difference of the number of rises between the winner and the loser emitted during the dark phase almost perfectly predicts the winner (AUC = 99.9 %, figs. 3 B). Initially, discrimination performance exponentially increases from chance level to maximum discrimination starting at 5 min after the beginning of a trial with a time constant of about 5 min (fig. 3 C). The prediction level of 94 % based on the physical factors (fig. S-3) is clearly surpassed after about 20 min. In that time the losing fish emitted on average 7.0 ± 5.7 rises and the winner just 1.2 ± 1.2.

The losing fish kept emitting higher numbers of rises than the winning fish throughout the dark phase of trials (figs. 3 D). In none of the trials did the competitors switch this behavior (figs. 3 E). The few instances were the fractions of rise counts of losing fish fall below 50 % are time windows with very few numbers of rises. Despite the ongoing emission of rises by the losing fish they failed to win the superior shelter.

Because of the low numbers of rises produced during the day and by winning fish, we focus in the following on rises produced by losing fish during the night. As detailed in the following, the number of rises produced by losing fish were dependent on the competitor’s sex, their physical differences, and the number of trials the fish already participated in. In contrast, we found no such dependencies for the number of rises emitted by winning fish.

### Loser against females emit more rises

In trials won by males, the losing competitor of either sex produced less rises than in trials won by females (*U* = 84.0, *p* = 0.02; fig. 4 A). Consequently, the number of rises produced by losing fish correlated positively with the difference in EODfs, because of the sexually dimorphic EODfs in *A. leptorhynchus* (*r* = 0.32, *p* = 0.05; fig. 4 B). Interestingly, in trials won by males the sex of the losing fish did not have an effect (*U* = 37.0, *p* = 0.36.0), whereas in trials won by females losing males produced more rises than losing females (*U* = 10.0, *p* = 0.036).

**Figure 4:**
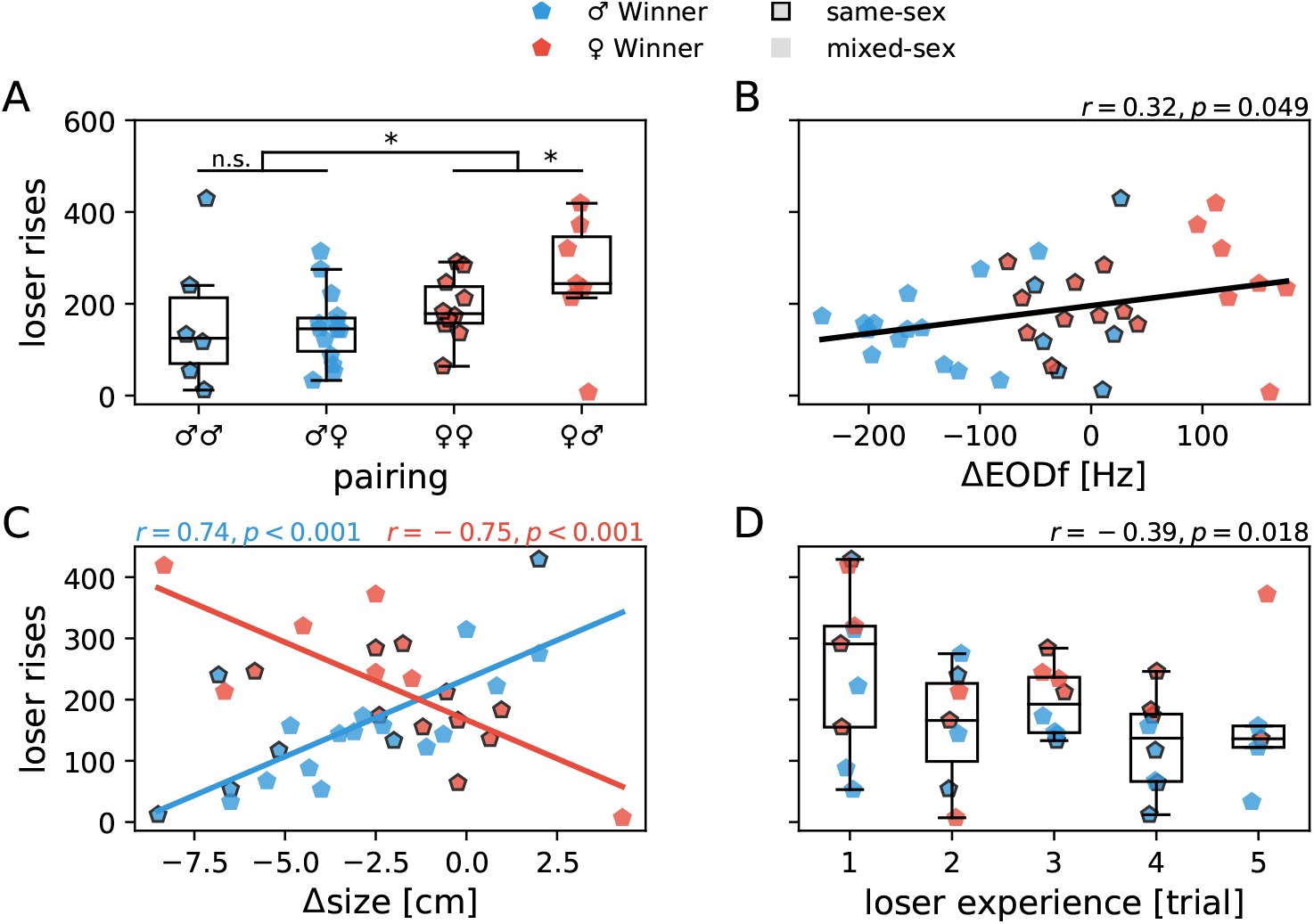
Dependence of rise counts of *losing* fish on physical differences between competitors and experience in the experiment. **A** Number of rises emitted by losers in different sex pairings. Markers indicate sex of winning opponent, first symbol in pairing category indicates sex of winner. In trials won by females (red) slightly more rises of the losing fish were observed than in trials won by males (blue). **B** With increasing difference in EODf to the winning fish (ΔEODf = EODf_loser_ − EODf_winner_), losing fish produced more rises. This effect rests on losing males emitting on average more rises than losing females in mixed-sex competitions (panel A) and males having higher EODfs than females. **C** In trials won by males (blue), the smaller the size difference to the winner (Δsize = size_loser_ − size_winner_) the more rises were produced by the losing opponent of either sex. In trials won by females (red), the opposite effect was observed. This effect can however also result from overall higher rise rates and larger size differences of males compared to females when losing against females (panel A). Correlation coefficients and their significance are displayed in corresponding colors. **D** With increasing experience in the experiments, losing fish produced less rises per trial.

### Sex-specific dependence of rise emission on body-size

The dependence on sex of the winner was even stronger when considering the contestants’ body-size. In pairings won by males, the number of rises emitted by the losers tended to increase with loser size (*r* = 0.42, *p* = 0.064) and decrease with winner size (*r* = − 0.51, *p* = 0.022). When regarding the difference in body-size, the number of rises emitted by the loser increased with the size of the losing fish as it approached and exceeded the size of the winning male (*r* = 0.74, *p* < 0.001, fig. 4 C). Backward elimination in a multiple linear regression model of number of rises in dependence on absolute sizes of competitors and their difference resulted in size difference as the only parameter (*t* = 4.72, *p* < 0.001) remaining in the significant model (*F*(1, 18) = 22.3, *p* < 0.001) with *R*^2^ = 0.55.

In pairings won by females, the number of rises emitted by losers decreased with loser size (*r* = −0.55, *p* = 0.023) and was unaffected be winner size. When regarding the difference in body-size, the number of detected loser rises decreased the more similar the competitors were in size (*r* = −0.75, *p* < 0.001, fig. 4 C), i.e. the effect was opposite to the one found for trails won by males. In a multiple linear regression model for trials won by females, size difference was the only parameter (*t* = −4.36, *p* = 0.001) remaining in the significant model (*F*(1, 15) = 18.98, *p* < 0.001) with *R*^2^ = 0.56 after backward elimination.

### Habituation of rise rates and loser effects

The number of rises produced by losers was independent from the outcome of preceding competitions, rejecting a loser effect on the communication behavior of *A. leptorhynchus*. However, the fish’s total experience in the experiment influenced the number of emitted rises. With increasing experience, i.e. the more trials a fish participated in, the number of rises produced by losing fish decreased (*r* = −0.39, *p* = 0.02; fig. 4 B). This habituation was independent of the paired sexes.

### Agonistic interactions

We detected in total 2 480 chasing behaviors and 804 agonistic interactions involving physical contact in the 19 trials were we recorded and evaluated these behaviors with infrared video. Agonistic interactions were exclusively detected during the dark-phase of each trial and stopped with or shortly after the onset of the light-phase (fig. 1 E).

In random visual inspections of video recordings we found agonistic behaviors to always be initiated by those fish later identified as winners. Per trial we observed on average 128 ± 72 chasing behaviors lasting each 7.4 ± 6.5 seconds and in addition 36 ± 21 physical contacts. The number of physical contacts between the competitors tended to increase with the number of chasings (*r* = 0.37, *p* = 0.12). Interestingly, none of the so far discussed factors had an impact on the number of both categories of agonistic interactions, including the competitor’s sexes, their size and EODf differences, and their experience in the experiment. In particular and similar to the rises, the number of interactions per trial were highly variable (contacts: *CV* = 0.55, chasings: *CV* = 0.58) and neither the number of contacts (*r* = −0.27, *p* = 0.24) nor the number of chasings (*r* = 0.30, *p* = 0.20) correlated with the number of rises.

### Sex-specific dependence of the duration of chasing behaviors on body size differences

However, the duration of the chasing behaviors were sensitive to these factors. The median chasing durations were shorter in male-male competitions compared to other pairings (*U* = 5, *p* = 0.003). In other pairings the median chasing durations were indistinguishable (fig. 5 A).

**Figure 5:**
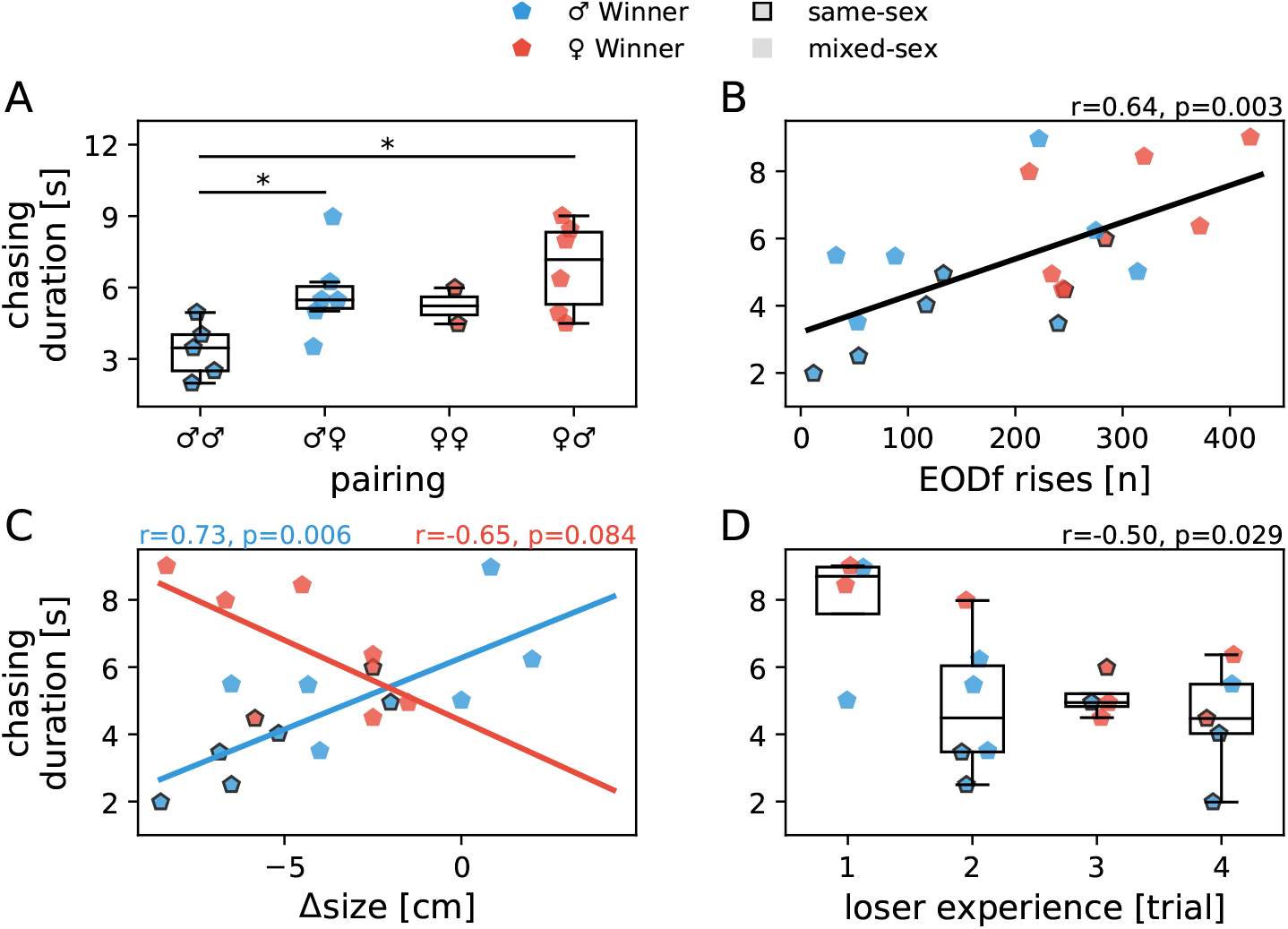
Agonistic interactions. While the number of agonistic interactions was independent of the physical characteristics of the opponents, the duration of chasing events showed similar dependencies as the number of rises. **A** The median chasing duration were the shortest in male-male interactions. In the annotations of the sex pairings the first symbol indicates the winner. **B** The median chasing duration increased with the number of rises emitted by losers. **C** In trials won by males the median chasing duration increased with losers approaching and exceeding the size of the winner (Δsize = size_loser_ − size_winner_). In trials won by females the opposite effect was observed, however similar explanations as for the quantity of rises in the respective pairings apply (fig. 4 B). Correlation coefficients and their significance are shown in corresponding colors. **D** With increasing experience in the experiments the median chasing duration decreased.

In trials won by males the median chasing duration tended to decrease with winner size (*r* = −058, *p* = 0.06) and was unaffected by loser size. However, it increased with the size of the losing fish relative to the size of the winner (*r* = 0.73, *p* = 0.006, fig. 5 C). Size difference remained after backward elimination the only parameter (*t* = 3.54, *p* = 0.006) in a linear regression model predicting median chasing duration (*F*(1, 9) = 12.5, *p* = 0.006) with *R*^2^ = 0.58.

In trials won by females median chasing duration was independent from absolute sizes but tended to decrease with decreasing size difference between competitors (*r* = −0.65, *p* = 0.08, fig. 5 D). Size difference was the last size parameter excluded by backward elimination, but did was not significant (*t* = −2.07, *p* = 0.084, *R*^2^ = 0.42).

Chasing durations were not correlated with differences in EODf between the competitors (*r* = 0.18, *p* = 0.45).

### Habituation of the duration of chasing behaviors

Beyond physical differences, the experience of the competitors in the experiment influenced the duration of chasing behaviors. The duration of chasings decreased with the number of trials the losing fish had experience with (*r* = −0.50, *p* = 0.03, fig. 5 E). The number of chasing behaviors was unaffected by experience (*r* = 0.05, *p* = 0.83). Finally, the observed communication behavior had an impact on the duration of chasings. In trials where losers produced more rises, chasings lasted longer (*r* = 0.64, *p* = 0.003, fig. 5 F).

### Some rises triggered agonistic interactions

Within about five seconds prior to agonistic interactions, rise rates accumulated over all agonistic interactions were increased (fig. 6 A & B). Because the baseline rate of rises was just one rise per minute, this does *not* imply that a burst of rises triggered agonistic interactions. Rather, the probability of a single rise to evoke an agonistic interaction was increased. In particular, chances for agonistic contacts were higher 0.7 s after a rise and chances for chasing behaviors were higher 1.6 s after a rise. Consequently, the fraction of rises occurring within five seconds prior to agonistic onsets exceeded the fraction expected from the corresponding times, the number of agonistic interactions times five seconds (agonistic contacts: 3.6 % versus 1.7 %, *t* = 2.9, *p* = 0.01; chasing behaviors: 9.6 % versus 5.7 %, *t* = −3.1, *p* = 0.007; fig. 6 B).

**Figure 6:**
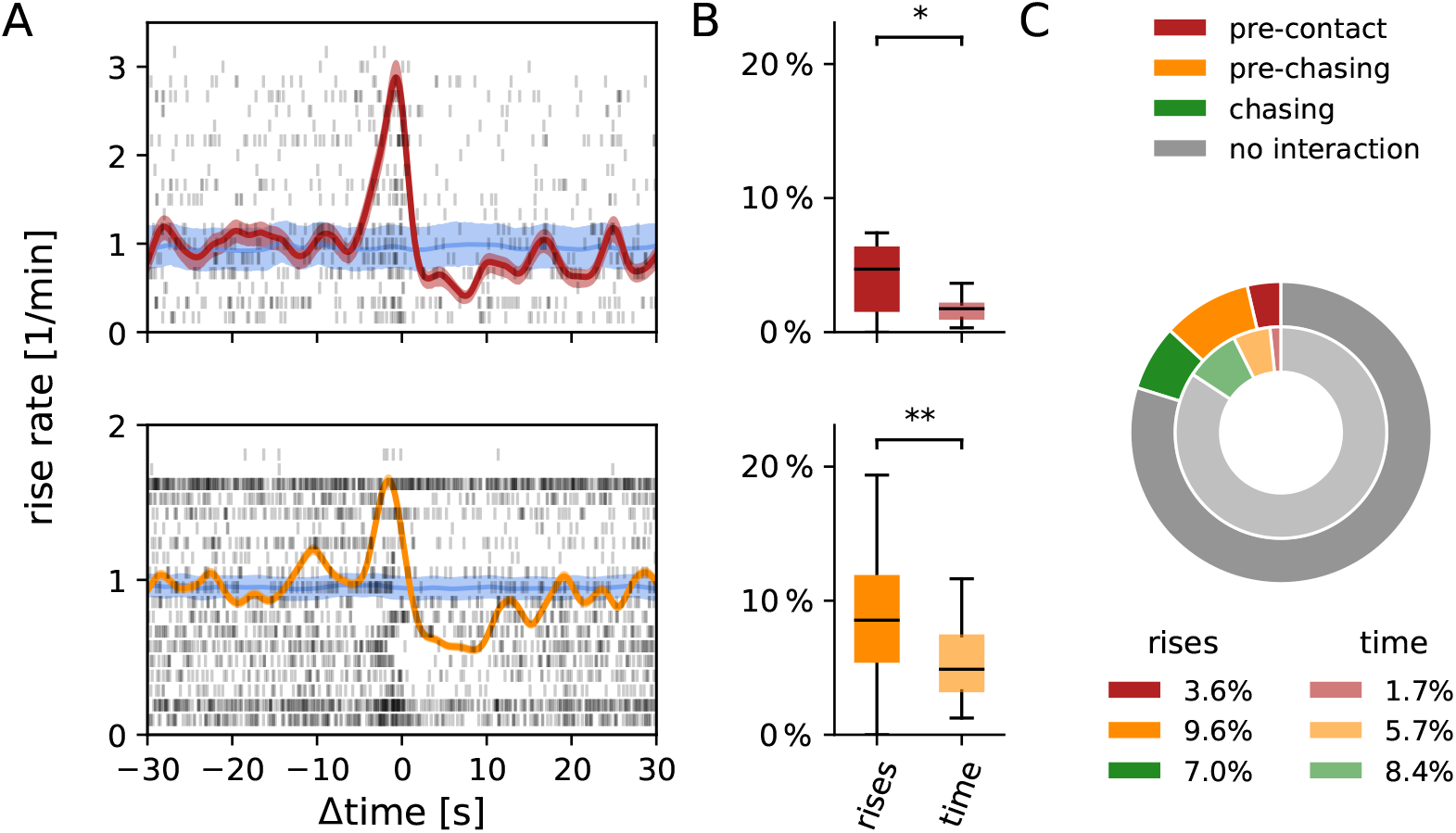
Rises trigger other fish to initiate agonistic attacks. **A** Rise times and rates relative to physical contacts (top) and the onset of chasing events (bottom). Each line of the rasters (black) shows rise times aligned to all agonistic events of a single competition trial. Corresponding rise rates (red/orange) were obtained by convolving rise times with Gaussian kernels with standard deviation of 1 s and normalizing by the number of agonistic events and trials. 98 % confidence intervals (reddish/orangish areas) were estimated by jackknifing. The Null-hypothesis (mean in blue, 1st and 99th percentile in bluish) was obtained by randomly permuting rise intervals. **B** Fraction of all rises within the dark phase of a trial that occurred within five seconds prior to agonistic contacts (top) or the onset of chasing behaviors (bottom) and the corresponding times these five-second windows make up relative to the total dark-phase lasting three hours. **C** Mean fractions of rises (outer circle) occurring within five seconds prior to physical contacts (red), prior to the onset of chasing behaviors (orange), and during chasings (green) and the corresponding times (inner circle) over all trials, indicating that in most cases rises did not evoke agonistic interactions.

### Reduced rise rates during chasings

Rise rates were approximately halved during chasings (fig. 6 A, bottom). After the onset of chasings rise rate was reduced for about ten seconds, clearly outlasting the average duration of chasing behaviors (7.4 s). Again this just implies that the chances to observe a single rise during chasing is reduced. About 7.0 % of rises occurred during chasing events, less than expected from the corresponding time covered by the chasing events (8.4 %, *t* = 3.4, *p* = 0.003, fig. 6 C).

### Most rises did not trigger agonistic interactions

All the rises emitted within five seconds prior to agonistic contacts and chasing events as well as during the chasings make up 20 % of all rises (fig. 6 C). Although this is disproportional more than expected from the corresponding times, the majority of the rises (80 %) did not trigger any obvious interaction. Also note that the fish engaged in actual agonistic interactions in form of chasing behaviors in just 8.4 % of the time. Vice versa, agonistic interactions were *not* exclusively triggered by rises emitted by losers. Rises preceded 15.6 % of all chasings and 19.4 % of physical contacts, respectively.

## Discussion

In staged competition experiments between pairs of the electric fish *A. leptorhynchus* of either sex we recorded electrocommunication signals, so called rises, as well as agonistic behaviors. Losers were characterized by relatively smaller body-sizes and by their continuously higher rise emission rates. Rises were costly for losers in that they raised chances to be attacked and to be chased for a longer time by winners. Our results suggest rises to signal an individual’s motivation to continue opponent assessment by stimulating ritualized fighting behaviors.

### Body-size as a proxy for RHP in A. leptorhynchus

An animal’s ability to win a fight, its RHP, is often correlated to body-size or weight, because physical strength is usually directly related to size (Parker, 1974; Archer, 1988; Arnott and Elwood, 2009). Accordingly, difference in body-size also was the best predictor for the outcome of competitions in our experiments (figs. 2 & S-3). Differences in motivation between individuals and sexes also influence the outcome of competitions (Enquist and Leimar, 1987; Arnott and Elwood, 2008; Dunham, 2008). In males an increased motivation and likelihood to compete has often been observed (Archer, 1988). In our experiments, males won competitions slightly more often than females, but mainly because of a bias of males being larger than females in these trials (fig. 2). Overall, average body size was independent of sex (fig. S-1 A). When evaluated separately, however, the correlation between body size and winning the superior shelter was more pronounced in trials won by males than trials won by females (fig. 2 B – E). This could reflect a higher valuation of suitable shelters by males compared to females (Arnott and Elwood, 2008).

### Relevant signals and interactions of A. leptorhynchus during competition

In the context of animal contests, individuals assess their own and/or the opponents RHP using cues, often including actively emitted signals, and adapt their behavior accordingly (Clutton-Brock et al., 1979; Enquist et al., 1990; Payne, 1998; Arnott and Elwood, 2009; Fernald, 2014). Properties of the continuously emitted electric field of *A. leptorhynchus*, in particular EOD frequency, could already be utilized in opponent assessment. Males increase both EODf and the androgen 11-ketotestosterone following environmental cues simulating the onset of breeding season (Cuddy et al., 2012) and in anecdotal observations males with higher EODf fertilize more eggs (Hagedorn and Heiligenberg, 1985; Henninger et al., 2018).

Outside the breeding season males with higher EODf have been found to be more territorial during the day (Raab et al., 2019) and to occupy their preferred shelter alone (Dunlap and Oliveri, 2002). Whereas in the latter study EODf was weakly but significantly correlated with body size, we did not have such a correlation (fig. S-1 A). Consequently, in our experiments predictive power of EODf on competition outcome was insignificant (fig. S-2), demonstrating only a minor role for EODf signaling RHP in addition to body size. However, further investigation is advisable, especially since attributes of EODs have been shown to influence competitions across electric fish species (Westby and Box, 1970; Dunlap and Oliveri, 2002; Cuddy et al., 2012)

The RHP of contestants usually affects their fighting behavior, e.g. quantity, intensity, duration, or point of giving up (Arnott and Elwood, 2009; Enquist et al., 1990; Taylor et al., 2001; Briffa and Elwood, 2004). While in *A. leptorhynchus* the number of agonistic interactions did not correlate with any of the measured parameters, the duration of chasings was dependent on the contestant’s difference in body-size, the main factor determining RHP, and the winner’s sex (discussed below). Interestingly, the number of rises emitted by losers were correlated in similar ways to the contestant’s RHP. This similarity together with the observation of rises frequently triggering agonistic attacks suggests rises to play a potential role in assessing the opponent.

### Electrocommunication with rises

*A. leptorhynchus* has been shown to use a rich repertoire of electrocommunication signals in social interactions (Smith, 2013; Benda, 2020). Some of the various types of chirps, transient elevations of EODf within less than about 500 ms, are used in agonistic, same sex encounters, to deter agonistic attacks (Hupé and Lewis, 2008; Henninger et al., 2018) and in courtship where synchronization of spawning is probably only one of their many functions (Hagedorn and Heiligenberg, 1985; Triefenbach and Zakon, 2003; Cuddy et al., 2012; Henninger et al., 2018). In contrast, evidence for the function of rises is scarce and inconsistent. Rises are characterized by smaller but much longer increases in EODf in comparison to chirps (Hopkins, 1974; Hagedorn and Heiligenberg, 1985). They vary considerably in their size (a few up to several tens of Hertz) and over three orders of magnitude in their duration (less than a second to up to a few minutes, Tallarovic and Zakon, 2002). The large number of rises we detected in our experiments clearly formed a continuous distribution of sizes and durations and we found no indication of distinct functional roles of rises of different sizes (Triefenbach and Zakon, 2008), refuting earlier attempts to categorize rises (Hagedorn and Heiligenberg, 1985; Tallarovic and Zakon, 2002; Dye, 1987).

At time scales longer than many seconds electric fish produce another category of signals where EODf stays at an elevated or reduced EODf (jamming avoidance responses: Bullock et al., 1972; Fortune et al., 2020, non-selective responses: Dye, 1987; Triefenbach and Zakon, 2008; Benda, 2020). If at all we observed only a few of such signals, but they appeared more like merged rises.

### Function of rises

Rises have been observed to be followed by attacks or bouts of aggression, both in *Eigenmannia* (Hopkins, 1974) and *A. leptorhynchus* (Triefenbach and Zakon, 2008), and to primarily be emitted by subordinates in *Apteronotus albifrons* (Serrano-Fernández, 2003). We also found rises in *A. leptorhynchus* to be primarily emitted by losing fish (fig. 3 A & B) and to trigger winners into initiating agonistic attacks within a few seconds (fig. 6). These common findings further support the hypothesis of rises being conserved signals in gymnotiform electric fish (Turner et al., 2007). In addition, chasings lasted longer the more rises the losers produced (fig. 5 B), and the number of agonistic behaviors was not reduced by more rises. This all contradicts the interpretation of rises as submissive signals (Hopkins, 1974; Serrano-Fernández, 2003) and as an general expression of stress (Smith, 2013), since stress levels are usually increased during agonistic encounters (Creel et al., 1996). Communication signals in general aim to alter the behavior of a receiver in a net beneficial fashion for the sender and they are only produced when the potential benefits outweigh the costs (Endler, 1993; Seyfarth et al., 2010). In animal contests they can convey information about physical condition and RHP (Davies and Halliday, 1978; Clutton-Brock et al., 1979), social status (Huyghe et al., 2005; Fernald, 2014), and motivation or behavioral intent (e.g. aggression: Triefenbach and Zakon, 2008; Kareklas et al., 2019, or submission: Hupé and Lewis, 2008; Batista et al., 2012) and often already convey sufficient information to settle competitions without the necessity of escalating costly fights (Arnott and Elwood, 2009). In our experiments winners could reliably be predicted within the initial 25 min of each trial based on the numbers of rises emitted by either fish (fig. 3 C). Nevertheless, losers continued to emit more rises than the winner until the end of the dark-phase (fig. 3 D). Never did we observe a switch in communication behavior between contestants (fig. 3 E). Therefore, rises were apparently not used to ultimately win competitions. What then is the purpose of rises?

### Sexual dimorphic behavioral traits

Males have been shown to be more territorial (Dunlap and Oliveri, 2002), can be assumed to be more aggressive (Zupanc and Maler, 1993), and show more intense dominance displays (Raab et al., 2019). Females, on the other hand, are known to be less territorial and more tolerant to the presence of con-specifics (Zupanc and Maler, 1993; Dunlap, 2002; Cuddy et al., 2012). All these observation suggest an increased resource valuation in males (territoriality at shelters) in comparison to females, and males being more motivated to win competitions. Our data further support this hypothesis, as discussed in the following.

In trials won by males, both the number of rises and the duration of chasing events increased with decreasing size difference between contestants (figs. 4 C and 5 C). The enhancement of both behaviors with decreasing size difference is not unusual, since individuals across species are more likely to compete with competitors of similar size, because of increased chances of success (e.g. Clutton-Brock et al., 1979; Arnott and Elwood, 2009; Enquist et al., 1990). A higher motivation of males could be perceived by losers by means of behavioral cues, interpreted as potentially higher costs when engaging in competition, and thus decreases the motivation of losers to compete. This could explain the overall lower rise production of losers in trials won by males (fig. 4 A) and the resulting shorter chasing events (fig. 5 A).

In trials won by females, we found opposite relations. With decreasing size difference less rises were produced by the losing fish and chasing duration decreased (figs. 4 C and 5 C). These negative correlations, however, are mainly carried by the sex of the losing fish. Males losing against females tend to be much smaller (fig. 2 E) and at the same time emit more rises (fig. 4 A) and interact longer during chasings (fig. 5 A) compared to all other pairings. Females competing against females were more similar in size (figs. S-1 A and 2 C), emitted less rises (fig. 4 A), and had shorter chasings (fig. 5 A). The higher intrinsic motivation of males in addition to the lower potential costs in competing with less territorial females could explain the enhanced rise production in males regardless of RHP of female opponents. In summary, the dependence of rise production by losing fish on sex demonstrates sexually dimorphic behavioral traits in *A. leptorhynchus*, arising from a higher motivation of males to win competitions despite substantial differences in RHP.

### Mutual assessment in A. leptorhynchus

Analysing the dependence of competition behaviors on the contestants’ RHP allows to differentiate between assessment strategies, i.e. pure-self, cumulative, or mutual assessment (Taylor et al., 2001; Payne, 1998; Enquist et al., 1990; Arnott and Elwood, 2009). In cumulative assessment costs arising from own actions during competitions and those being inflicted by opponents are accumulated until an endurance threshold is reached and the animal retreats (Payne, 1998). In our experiments, however, this threshold never seems to be reached, because rise emission and agonisitic interactions went on until being terminated by the lights being switched on and fish becoming inactive. For pure self assessment a positive correlation between both contestants absolute RHP and the extend of behaviors associated with competition is expected (Arnott and Elwood, 2009). This can be rejected, because absolute body-size of either winner or loser did not predict competition outcome (fig. 2 F), and both quantity of rises and duration of chasing events rather decreased with winner size.

Both, communication and agonistic behaviors remained steady throughout single trials. They can be assumed to be low costs behaviors, because no injuries or other forms of negative consequences resulted from them. This supports mutual assessment where low cost competition behaviors are repetitively performed in order to accurately assess an opponent’s RHP relative to the own (Enquist et al., 1990; Clutton-Brock et al., 1979). Furthermore, animals are expected to improve in accuracy of assessing the opponent with increasing experience (Enquist et al., 1990; Grosenick et al., 2007). This matches previous observations on decreasing competition intensity over trials in another gymnotiform electric fish (Westby and Box, 1970) as well as our own observations on *A. leptorhynchus* where the number of emitted rises and the duration of chasing events decrease with the fish’s experience in the competition experiment (figs. 4 D and 5 D).

### Dominance in A. leptorhynchus

Competitions *per se* are not exclusively used to directly secure access to resources, but also to establish dominance hierarchies, which indirectly regulates access to resources (Janson, 1985; Wauters and Dhondt, 1992; Sapolsky, 2005; Taves et al., 2009). After establishing dominance, knowledge about an individual’s social status and the interpretation of cues and signals indicating it can prevent costly repetitive fighting and therefore be beneficial for all individuals involved (Fernald, 2014; Huyghe et al., 2005). Characteristics of social hierarchies and behavioral correlates of dominance vary widely across species (Clutton-Brock et al., 1979; Cigliano, 1993; Sapolsky, 2005; Strandburg-Peshkin et al., 2018). In group living species, beyond regulating access to resources, social hierarchies often come along with complex social dynamics, like, for example, leader-follower dynamics (e.g. Strandburg-Peshkin et al., 2018; Janson, 1990). In solitary species, on the other hand, dominance is primarily associated with resource-based benefits (Cigliano, 1993).

Dominance hierarchies have also been suggested for *A. leptorhynchus* (Hagedorn and Heiligenberg, 1985; Dunlap and Oliveri, 2002). Since behavioral observations suggests *A. leptorhynchus* to be a solitary living species (Stamper et al., 2010; Raab et al., 2019; Henninger et al., 2020), this dominance can be assumed to be mainly resource based. In appropriate habitats with suitable shelters, *A. leptorhynchus* is quite abundant, but the fish distribute themselves independently from each other and do not form large social groups (Stamper et al., 2010; Raab et al., 2019, own observations in Colombia 2016, 2019). Nevertheless, the high abundance in their natural habitats (Albert and Crampton, 2005) require these fish to frequently interact with con-specifics. Thus, repetitive interactions are likely and the establishment of dominance can be assumed to be advantageous.

Previous studies suggest male but not female *A. leptorhynchus* to form a dominance hierarchy (Hagedorn and Heiligenberg, 1985; Dunlap and Oliveri, 2002). Indeed, as discussed above, males seemed to be more motivated to win competitions. Nevertheless, competition outcome was independent from the contestants’ sex and mainly determined by their relative body-size (fig. S-3). Dominance in *A. leptorhynchus* thus appears to be sex-independent, in line with similar studies on other gymnotiform electric fish (Batista et al., 2012; Zubizarreta et al., 2020).

### Rises in the social hierarchy of A. leptorhynchus

The behaviors of animals, including the emission of communication signals aim at benefits for a focal individual (Parker, 1974; Seyfarth et al., 2010). In our experiments, however, the continuous emission of rises did not result in direct, conspicuous benefits for losers, but rather increased costs in terms of enhanced agonistic attacks towards them. Why keep the fish emitting rises?

In social hierarchies dominants often use agonistic attacks to keep subordinates under control (Clutton-Brock et al., 1979; Creel et al., 1996; Janson, 1985; Sapolsky, 2005). In *A. leptorhynchus* subordinates could emit rises to signal their motivation to continue assessment, with the aim to reduce relative dominance (e.g. Kareklas et al., 2019). Dominant fish, on the other hand, counteract with agonistic attacks (e.g. Creel et al., 1996; Janson, 1985; Clutton-Brock et al., 1979). The interplay and balance between rises and agonistic attacks could define the relative dominance between contestants and regulate skewness in access to resources. This would imply that motivation of the fish in the competition depends on the valuation of not only present but also regularly encountered resources, most likely food, that were absent during our experiments.

This hypothesis on a possible benefit of continuous loser rise emission is supported by a single exceptional trial, where the dominant fish shared the superior shelter with the subordinate at the end of a trial. In this mixed-sex trial, the smaller male (11.9 cm) was continuously emitting 180 rises during the dark phase and apparently succeeded in reducing relative dominance difference to the larger female (12.5 cm, no rises during dark-phase) by gaining access to the shelter.

## Conclusion

Male *A. leptorhynchus* seem to be more motivated to win staged competitions for a superior shelter than females. Nevertheless, contest outcomes were mainly determined by relative body-size, reflecting their overall fighting ability, their RHP. During competition, *A. leptorhynchus* interact physically by means of ritualized fights and use rises as distinct electrocommunicaion signal. The way how the extent of both behaviors depends on the contestants fighting ability suggests *A. leptorhynchus* to assess their opponents during contests (mutual assessment). Here, rises are almost exclusively emitted by losers and seem to signal their motivation to continue physical assessment. Rises triggered agonistic attacks and enhanced the duration of chasing events. The motivation to continue assessment could reflect a loser’s attempt to reduce relative dominance, which is counteracted by the dominant fish with agonistic attacks.

## Acknowledgements

Supported by Deutsche Forschungsgemeinschaft via mini RTG “Sensory flow processing across modalities” by the excellence cluster “Center of Integrative Neuroscience” (CIN EXC 307).

## Author contributions

T.R. designed the experiment, measured and analyzed data, and wrote the manuscript. S.B. and S.E. measured the data. J.B. discussed the experiment, advised data analysis, and edited the manuscript.

## Supplemental figures

**Figure S-1:**
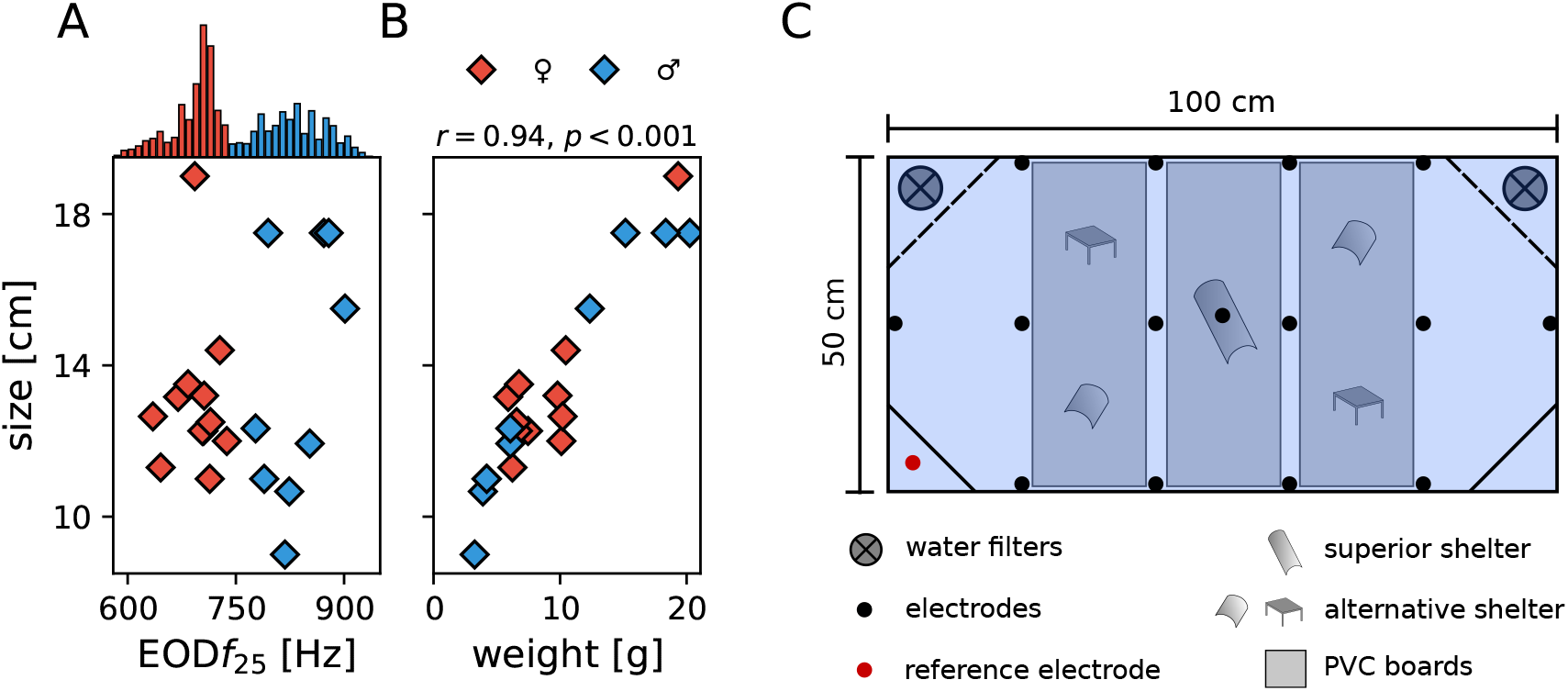
Physical characteristics of the fish and experimental setup. **A** EODf and size (length) of each of the 21 *A. leptorhynchus* participating in the competition experiments. EODf was not correlated with size, neither for all fish nor within the sexes. EODfs were temperature corrected to 25 °C. *A. leptorhynchus* males (blue) were identified by their higher EODfs compared to the EODf of females (red). The histogram on top shows the distribution of EODfs of either sex measured in all trials for every five minutes. **B** Weight and size of the fish were independent of sex and increase proportional to each other, i.e. fish grow mainly in length. **C** Setup for the staged competition experiments. The tank was equipped with one high quality shelter (center tube) and four low quality shelters (two short tubes and two tables) attached to PVC-boards to keep them in place. A total of 15 electrodes (black circles) were distributed in the tank to record electric behaviors of interacting fish. Two air-powered water-filters were placed in the corners behind PVC boards with netted windows (dashed lines). The other two corners were shielded with PVC boards (black lines), one of them contained the reference electrode (red circle).

**Figure S-2:**
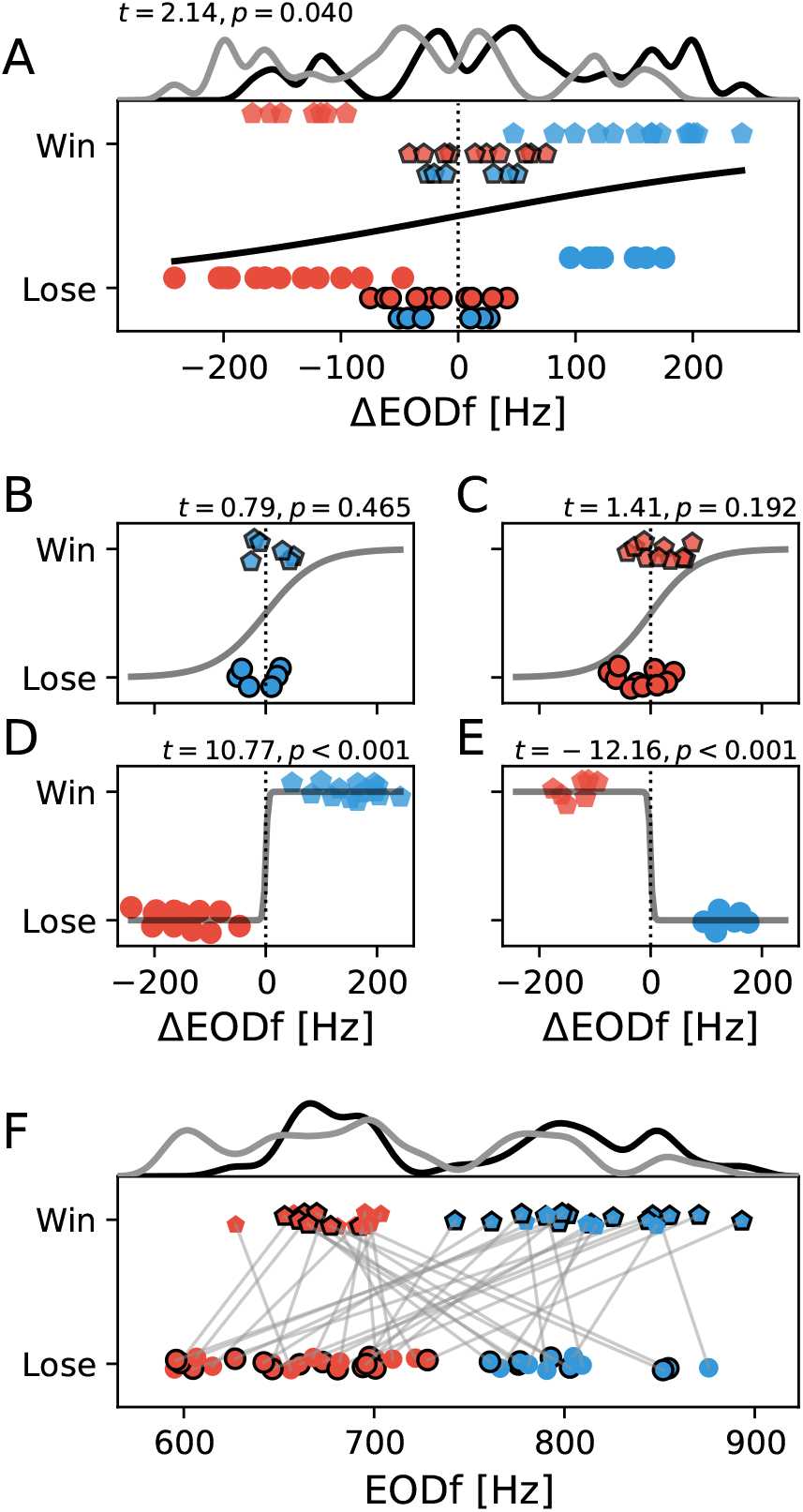
EOD frequencies of winners and losers. Same annotation scheme as in fig. 2. **A** Winners tend to have higher EODfs than their opponents. **B, C** Winners of same-sex encounters do not have significantly higher EODfs than losers (AUC=73 %). **D, E** In mixed sex competitions the sexual dimorphic EODf does not predict competition outcome. Males with their higher EODf can lose against females. More males (*n* = 14) were winning mixed-sex trials than females (*n* = 7), explaining the overall trend of winners having higher EODfs than losers (panel A) **F** Absolute EODf of winners and losers convey even less information about the outcome of the competitions (AUC=65 %).

**Figure S-3:**
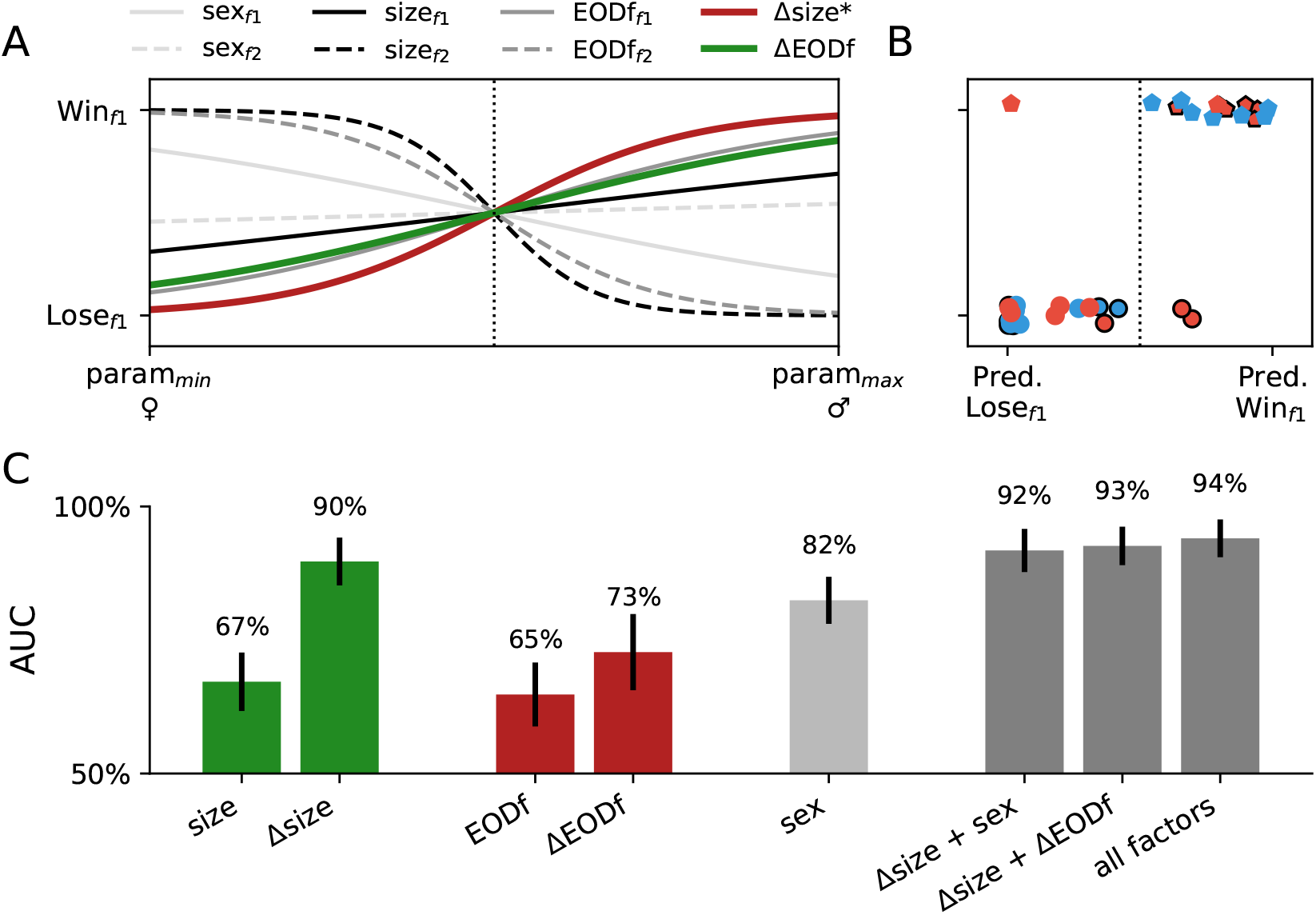
Logistic regression models predicting the outcome of the competition experiments. **A** Visual representation of the impact of all factors (sex, absolute size, and EODf of fish *f* 1 and *f* 2 competing with each other, as well as their differences in size and EODf, see legend) on the generalized linear mixed model with a logistic link function, corresponding to tab.1. The steeper the logistic functions the more discriminative the respective factor. Size difference, Δsize, is the only significant factor. **B** Predictions of the GLM on fish_1_ of each of the 37 competition trials being the winner or loser based on all factors shown in A. Same marker code as in fig. 2. **C** Area under the curve (AUC, with standard deviation estimated by bootstrapping) extracted from a receiver-operating-characteristic (ROC) analysis to quantify the performance of single factors and combinations of factors to discriminate winners of competitions from losers. Chance level is at AUC = 50 %.

## Notes

### Competing Interest Statement

The authors have declared no competing interest.

